# A panel of TDP-43-regulated splicing events verify loss of TDP-43 function in amyotrophic lateral sclerosis brain tissue

**DOI:** 10.1101/2023.02.03.527079

**Authors:** Maize C. Cao, Brigid Ryan, Jane Wu, Maurice Curtis, Richard Faull, Mike Dragunow, Emma L. Scotter

## Abstract

TDP-43 dysfunction is a molecular hallmark of amyotrophic lateral sclerosis (ALS) and frontotemporal dementia (FTD). A major hypothesis of TDP-43 dysfunction in disease is the loss of normal nuclear function, resulting in impaired RNA regulation and the emergence of cryptic exons. Cryptic exons and exon changes are emerging as promising markers of lost TDP-43 function in addition to revealing biological pathways involved in neurodegeneration in ALS/FTD. In this brief report, we identified markers of TDP-43 loss of function by depleting *TARDBP* from post-mortem human brain pericytes, a manipulable *in vitro* primary human brain cell model, and identifying differential exon usage events with bulk RNA-sequencing analysis. We present these data in an interactive database (https://www.scotterlab.auckland.ac.nz/research-themes/tdp43-lof-db-v2/) together with seven other TDP-43-depletion datasets we meta-analysed previously, for user analysis of differential expression and splicing signatures. Differential exon usage events that were validated by qPCR were then compiled into a ‘differential exon usage panel’ with other well-established TDP-43 loss-of-function exon markers. This differential exon usage panel was investigated in ALS and control motor cortex tissue to verify whether, and to what extent, TDP-43 loss of function occurs in ALS. We find that profiles of TDP-43-regulated cryptic exons and changed exon usage discriminate ALS brain tissue from controls, verifying TDP-43 loss of function as a pathomechanism in ALS. We propose that TDP-43-regulated splicing markers with most predictive value for therapeutic intervention will be those based on splicing events that occur both in tissues/biofluids amenable to sampling, and in brain tissue susceptible to disease.

## Introduction

Amyotrophic lateral sclerosis (ALS) and tau-negative frontotemporal dementia (FTD) represent the ends of a neurodegenerative disease spectrum characterized by TDP-43 proteinopathy (1-3). TDP-43 normally functions as an RNA-binding protein in the nucleus, but in disease its misfolding and mislocalisation to the cytoplasm is proposed to cause loss of nuclear function (4-6). Functions of TDP-43 include regulation of RNA expression levels, stability and splicing, and suppression of cryptic exons (7-11). Patterns of differential splicing and cryptic exon emergence with experimental TDP-43 depletion are more robust to cell type than patterns of differential gene expression (4), and have recently shown promise as markers of TDP-43 loss of function (LOF) in disease. However, for splicing changes to be used as peripheral/biofluid biomarkers to gauge cellular dysfunction and the efficacy of therapeutic interventions, i) those splicing changes must also occur in brain tissue and ii) loss of TDP-43 function must be a major driver of disease pathogenesis.

The normal physiological functions of TDP-43 involve the regulation of different stages of coding and non-coding RNA metabolism, both in the nucleus (including biogenesis and splicing) and cytoplasm (including stability and degradation) (7, 8). TDP-43 has two RNA recognition motifs (RRMs) that facilitate binding to RNA substrates, a nuclear localisation sequence (NLS) that allows nucleocytoplasmic shuttling and a glycine-rich and relatively unstructured C-terminus for protein-protein interactions (12). ALS-causing mutations in the *TARDBP* gene encoding TDP-43 are concentrated in the glycine-rich region, which may implicate altered protein-protein interaction in disease but more likely demonstrates that mutation of functional domains involved in nuclear shuttling and RNA binding is incompatible with life (13).

Nuclear TDP-43 regulates the expression and splicing of thousands of transcripts (4, 9), including regulatory non-coding RNAs (14) and mRNAs involved in nucleocytoplasmic transport, neurotransmitter release, mitochondrial motility and microtubule stability (9, 10, 15-19). Nuclear TDP-43 also autoregulates its own mRNA levels through binding its own 3’ UTR (20). Loss of functional TDP-43 from the nucleus would therefore dysregulate diverse cellular pathways, and further contribute to TDP-43 overproduction, phase separation/aggregation and sequestration (21-25). While TDP-43 loss from the nucleus had been described upon first discovery of ALS/FTD neurons with cytoplasmic TDP-43 aggregation (26, 27), given the dominant inheritance of *TARDBP*-linked ALS/FTD and the recapitulation of disease phenotypes by TDP-43 overexpression (28-31) the favored hypothesis at that time was a gain of toxic function of aggregates. More recently however, TDP-43 nuclear loss has been reported even in neurons without TDP-43 aggregation (32, 33) and disease phenotypes have also been recapitulated by TDP-43 knockdown (34-37), suggesting that TDP-43 LOF is a significant pathomechanism in ALS/FTD.

TDP-43 LOF has been difficult to verify in human ALS/FTD brain tissue. Differential expression of a range of genes has been demonstrated in human disease and various models but gene expression changes are difficult to validate as being a direct consequence of dysfunctional TDP-43. However, TDP-43 also regulates splicing, by suppressing or enhancing the expression of exons (8-10, 12) and changes in splicing are more specific than differential gene expression to TDP-43 LOF (4). Recently, the loss of TDP-43 function as a suppressor of cryptic exons has been highlighted in both TDP-43 knockdown models and in ALS/FTD, particularly in the genes *STMN2* and *UNC13A* (16-19). Cryptic exons (or pseudoexons) usually result from improper intron removal, and can arise through single nucleotide polymorphisms (SNPs) in acceptor/donor splice sites or the activity of spliceosome or RNA-binding proteins such as TDP-43 (38). Cryptic exons may alter the open reading frame and/or cause nonsense-mediated decay of the mRNA by introducing a premature stop codon, mimicking gene deletion (39). Thus, splicing events, and particularly cryptic exon emergence, are now being used as reporters of TDP-43 LOF. Identifying additional TDP-43-regulated splicing events would enable verification that TDP-43 LOF occurs in ALS/FTD and indicate pathways changed in disease, and these events could also serve as markers of TDP-43 dysfunction and its therapeutic reversal.

Intriguingly, cryptic exons that emerge in human models of TDP-43 LOF and in disease are generally not recapitulated in mouse models (4, 11, 40, 41). TDP-43 also has distinct sets of molecular targets within a given species but in different cell types (4, 11, 40, 41). The activity of TDP-43 therefore appears to be dependent on transcriptomic context, and human brain-derived transcriptomes are likely to be most suitable for identifying additional TDP-43 splicing events relevant to human ALS/FTD. Here, we examined TDP-43 LOF in primary human brain-derived pericyte cell lines from multiple control donors subjected to *TARDBP* depletion. These lines are genotypically heterogeneous, untransformed, and a manipulable primary human brain cell model for inducing TDP-43 LOF to identify splicing markers thereof. As non-neuronal cells they also expand the transcriptomic ‘search space’ beyond that of neuronal models used previously (5, 17, 19).

In this study, we identified cryptic exons caused by TDP-43 LOF by bulk RNA sequencing of human brain-derived pericytes with siRNA-mediated *TARDBP* depletion. We then examined the usage of our novel exon targets and previously reported exon targets in ALS brain tissue, to determine whether profiles of differential exon usage and cryptic exons could contribute to ALS/FTD pathomechanisms and thus be harnessed as peripheral biomarkers or targets for intervention.

## Results

### TARDBP silencing in a primary human brain pericyte model identifies known and novel TDP-43-regulated splicing events

Post-mortem human brain pericytes derived from the middle temporal gyrus (MTG) or motor cortex (MC) were isolated and treated with scrambled siRNA (scr) or siRNA against *TARDBP* (siTDP) to examine TDP-43 LOF. RNA samples were extracted from paired samples (scr and siTDP) from 12 donor cell lines (**Table S1**). Of these RNA samples, 9 pairs were sent for RNA-sequencing as reported previously (42) (accession number GSE223747) and 3 pairs (extracted from cell lines from different donors from those sent for sequencing) were used as a ‘replication cohort’ to validate RNA-sequencing results by qPCR. Successful *TARDBP* depletion from the RNA-sequencing cohort was demonstrated by differential gene expression analysis using the *DESeq2* package in R software (**Fig. 1A**, GEO GSE223747 Supplementary File 1), and from the replication cohort by qPCR for *TARDBP* (**Fig. 1B)** and immunocytochemistry for TDP-43 (**Fig. 1C**).

**Figure 1.**
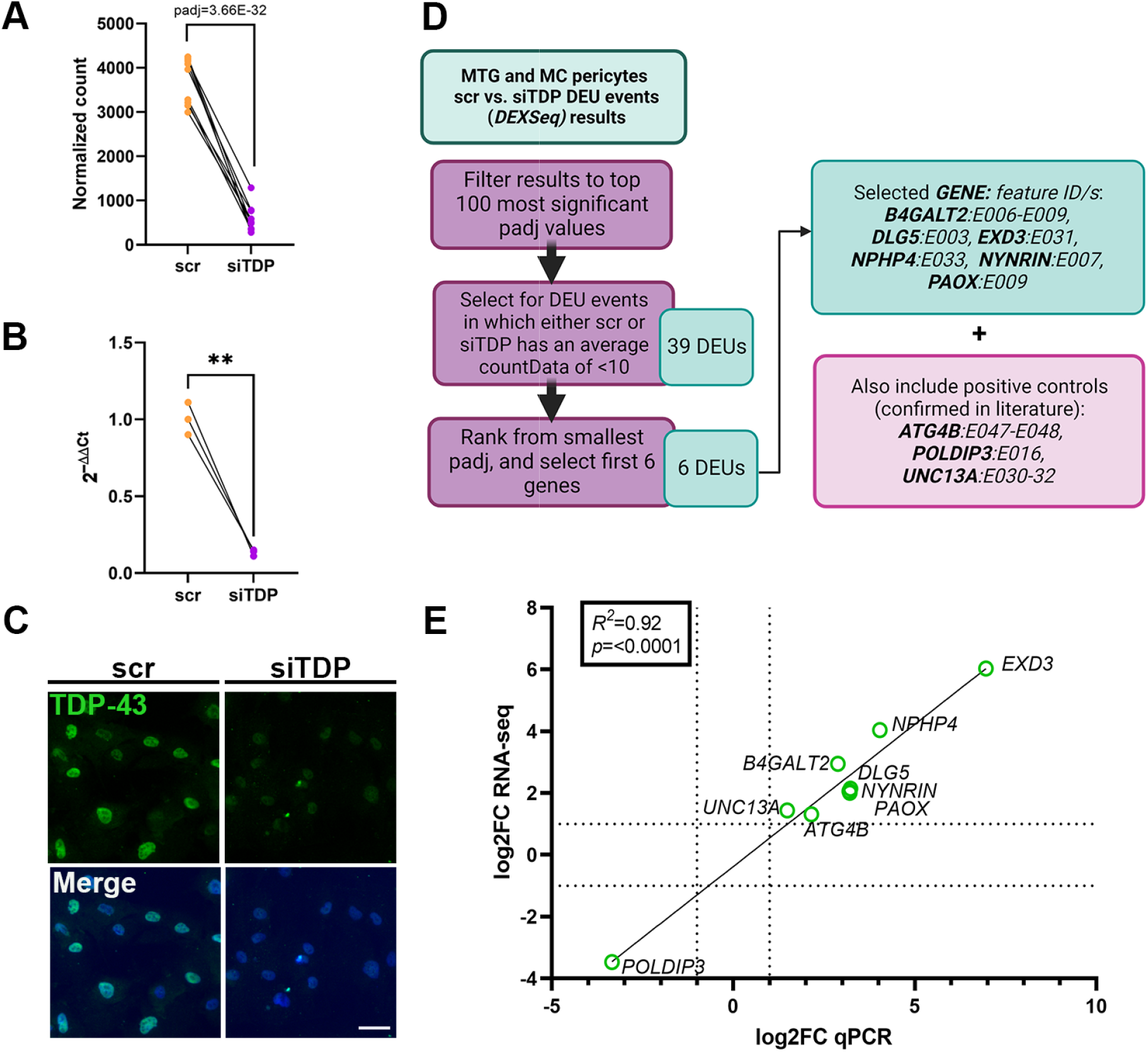
Differential exon usage in human brain pericytes with TDP-43 knockdown. A)*TARDBP* depletion from pericytes treated with scr or siTDP for 96 hours was confirmed in the RNA-seq cohort (n= 9) by RNA-Seq (paired Wald test with Benjamini-Hochberg correction (padj = 3.66E-32)), and B) in the replication cohort (n= 3) by qPCR (paired t-test: **, p < 0.005). C) Depletion of TDP-43 protein levels in pericytes treated as in A & B was confirmed by immunocytochemistry (TDP-43, green; Hoechst, blue). Representative donor cell line shown. Scale bar, 50 μm. D) Flow chart showing the selection of DEU events of interest from DEXSeq analysis to validate by qPCR. E) Correlation plot of log2FC determined by RNA-Seq and log2FC determined by qPCR of MTG and MC pericyte DEU events. All DEU events reached a log2FC of >1 or < -1 (shown as dotted lines).

Differential exon usage (DEU) analysis of the RNA-sequencing data was performed using the *DEXSeq* package in R software. This package quantifies the relative usage of exonic elements, which are exons or parts of exons (annotated by feature ID, e.g. E001) and can therefore detect exon exclusion and exon inclusion; cryptic exon and intron retention events; and differential usage of non-coding elements such as the 5’ and 3’ UTRs. After *TARDBP* knockdown in pericytes, DEU analysis identified differential usage of 1113 exonic elements in 564 genes including 3 previously published positive control DEU events in *ATG4B* (cryptic exon (11, 43)), *POLDIP3* (exon skipping (44)) and *UNC13A* (retained intron (16)) (GEO GSE223747 Supplementary File 2, padj < 0.05). These data can also be explored interactively online, and compared to seven other TDP-43-depletion datasets we meta-analysed previously (4): https://www.scotterlab.auckland.ac.nz/research-themes/tdp43-lof-db-v2/.

Of these 1113 DEU events, 6 were selected to validate, by qPCR of a replication cohort of pericyte lines with *TARDBP* knockdown. These 6 DEUs were selected because they had among the most significant padj values, and their exon expression (countData) either changed to, or from, very low expression level (<10 counts) with siTDP treatment (**Fig. 1D**). This would indicate that exonic element inclusion or exclusion was dependent on the presence of TDP-43. The 6 selected DEU events were in genes *B4GALT2, DLG5, EXD3, NPHP4, NYNRIN* and *PAOX*. Primers were designed to target the specific exon sequence that was changed. The 3 positive control DEU events in *ATG4B, POLDIP3* and *UNC13A* were also validated in directional change (i.e. suppression or inclusion) by qPCR.

For these 9 DEU events, log2 fold-changes (log2FCs) in the replication cohort by qPCR were strongly correlated with log2FCs in the RNA-sequencing cohort by RNA-seq (**Fig. 1E**, *R* = 0.9193). Exon coverage traces revealed exon or intron inclusion in *ATG4B, UNC13A, B4GALT2, DLG5, EXD3, NPHP4*, and *NYNRIN*; an exon variant in *PAOX* and exon skipping in *POLDIP3*. All 9 genes showed differential exon usage in the same direction (i.e. suppression or inclusion) by qPCR as by RNA-seq, with 6 genes meeting statistical significance (*ATG4B, POLDIP3, B4GALT2, EXD3, NPHP4* and *NYNRIN*) (p < 0.05) by qPCR (**Fig. 2**).

**Figure 2:**
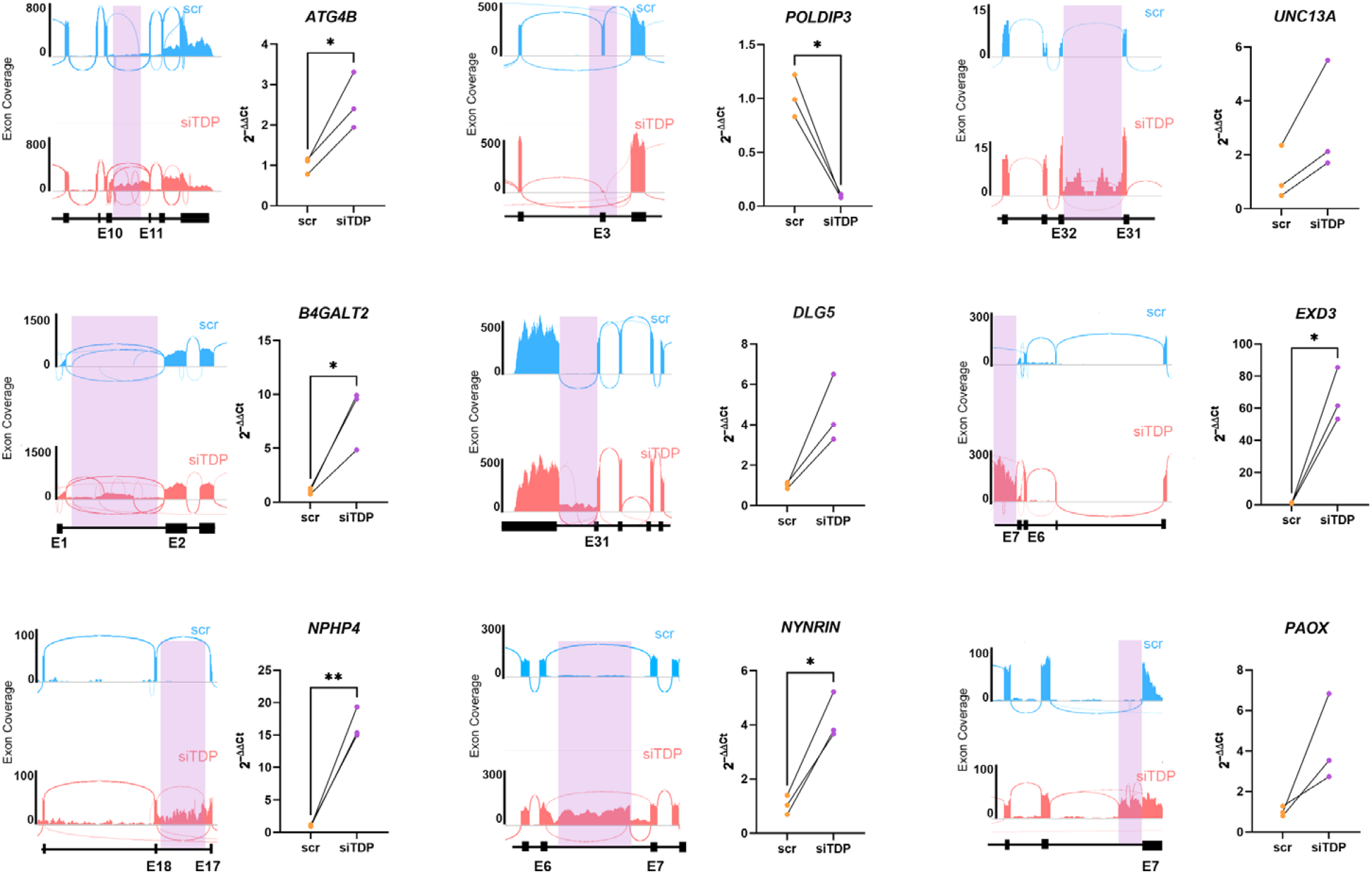
Validation of differential exon usage events in human brain pericytes with replication cohort. (*Left panels*) Exon coverage plots of DEU events (purple boxes), in positive control genes and genes identified in pericytes treated with scr (blue) or siTDP (red) for 96 hours. Exon-intron schematic 5’ (left) to 3’ (right) direction. (*Right panels*) Fold changes in the indicated DEU event in the replication cohort (n= 3) by qPCR showing the same direction of change between scr and siTDP as the RNA-seq cohort. *ATG4B, POLDIP3, B4GALT2, EXD3, NPHP4*, and *NYNRIN* were significantly changed (Paired t-test: *, p < 0.05; **, p < 0.01), and the other targets showed consistent direction and magnitude of change between biological replicates.

### Assembling a panel of known and novel TDP-43-regulated splicing events

Having identified and verified novel TDP-43-dependent splicing events in human brain pericytes (DEU events in *B4GALT2, DLG5, EXD3, NPHP4, NYNRIN*, and *PAOX*), we combined these into a ‘panel’ with DEUs from our previous TDP-43 LOF meta-analysis (4) (DEU events in *GOSR2, IL18BP, MARK3, MMAB, NDUFA5* and *USP31*) and positive controls previously tested in the literature (DEU events in *ATG4B, POLDIP3*, and *UNC13A* (as above), and additionally *STMN2* (17, 19)) (**Fig. 3A**). We hypothesized that this panel of validated TDP-43 LOF-induced DEU events could enable us to examine the extent of TDP-43 LOF in human brain tissue, and act as peripheral biomarkers.

**Figure 3.**
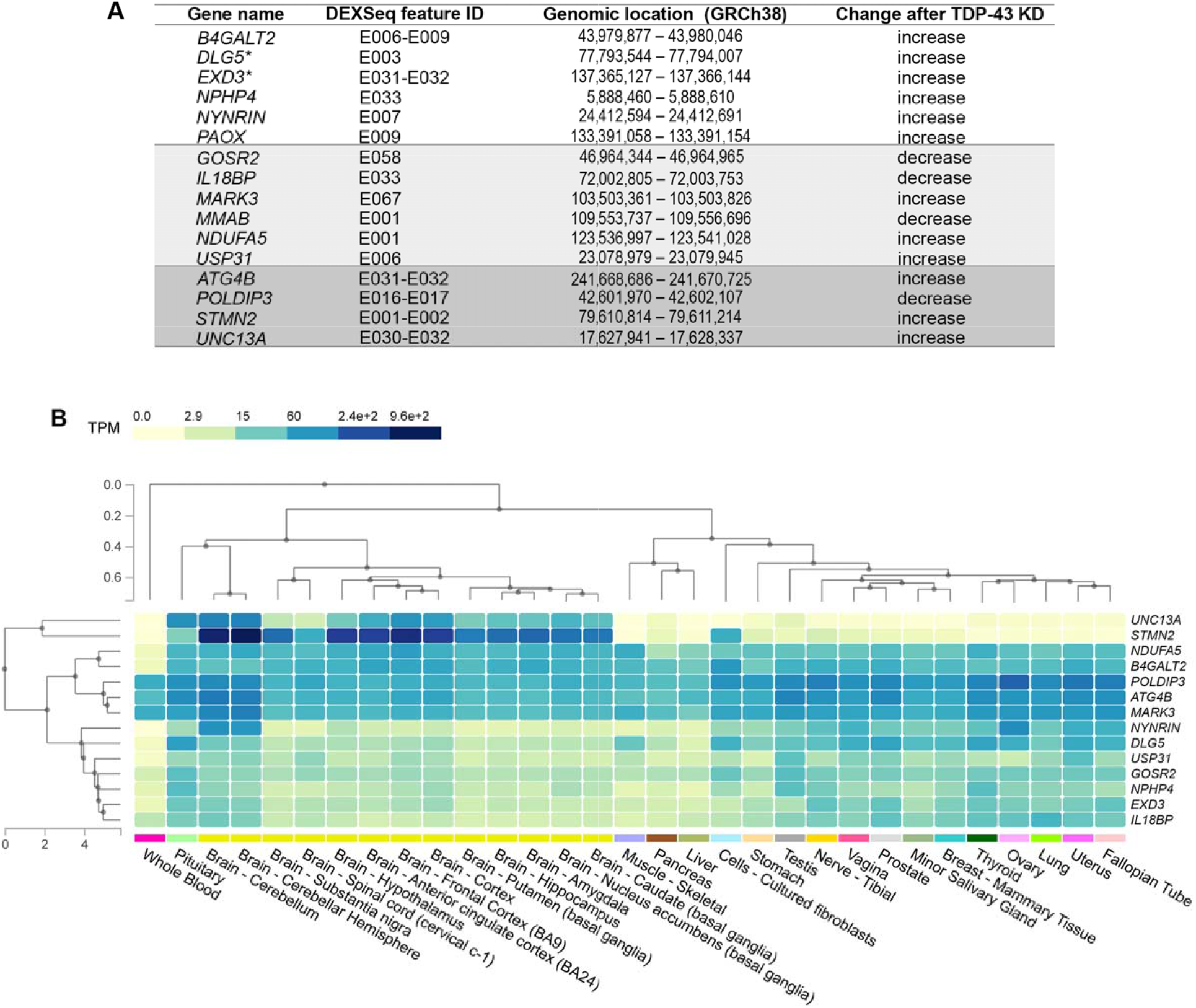
Differential exon usage marker panel as assembled from DEU events with TARDBP knockdown in human brain pericytes and other human-derived cells. A) Table showing panel of DEU events to test in human brain tissue. Non-shaded rows: DEU events identified with TDP-43 LOF in pericytes (this study); light-shaded rows: DEU events identified with TDP-43 LOF meta-analysis of various models (4); dark-shaded rows: DEU events previously reported by TDP-43 LOF studies. B) Heatmap showing expression of the DEU panel genes across well-studied tissues. Plot retrieved from Genotype-Tissue Expression (GTEx) Portal V8 (www.gtexportal.org).

We therefore next examined expression of the genes with DEU events across tissue types, including brain, blood and cultured fibroblasts, using publicly available data from the Genotype-Tissue Expression (GTEx) database (**Fig. 3B**). Genes with moderately to highly expressed transcripts in human brain are *UNC13A, STMN2, NDUFA5, B4GALT2, POLDIP3, ATG4B* and *MARK3*, while *NYNRIN* is moderately expressed in cerebellum but no other brain regions. Of these, *POLDIP3, ATG4B* and *MARK3* are also highly expressed in whole blood, with low expression of *NDUFA5* and *B4GALT2*, and no expression of *UNC13A* or *STMN2* in whole blood. Cultured fibroblasts express all genes in the panel except *UNC13A*.

### A panel of known and novel TDP-43-regulated splicing events discriminates ALS from control brain tissue, verifying TDP-43 loss of function in ALS

Fresh-frozen tissue samples from motor cortex of ten control (case IDs ‘H#’) and nine ALS cases (case IDs ‘MN#’) were obtained for RNA extraction, followed by cDNA synthesis and qPCR using primers to selectively amplify the DEU events. Age, sex and post-mortem delay were matched as closely as possible (**Table S2**).

The relative fold change (2^-ΔΔCt^ method) for ALS samples and controls for each DEU event is presented in **Fig. 4A**. All DEU events tested could be detected within 40 PCR cycles except for that in *MMAB*. Therefore, Ct data for *MMAB* could not be obtained, presumably because *MMAB* gene expression was too low. While 3 DEU events were expected to show a decrease with TDP-43 LOF (*MMAB, IL18BP* and *GOSR2*), the mean 2^-ΔΔCt^ value for ALS cases was increased or trended towards an increase for almost all DEU events compared to controls. However, for the DEU events in *DLG5, NDUFA5, USP31* and *STMN2*, which were predicted to increase, four cases in the ALS cohort (MN13, MN28, MN4 and MN9) demonstrated over 6-fold increases. Three of these DEU events (in *DLG5, NDUFA5* and *USP31*) were significantly increased in ALS tissue, as had been seen with TDP-43 LOF in pericyte RNA-seq data and/or meta-analysis of TDP-43 LOF models (4). Positive controls *ATG4B, POLDIP3, STMN2* and *UNC13A* did not show a significant change in ALS samples, possibly due to the mixed cell-type nature of brain tissue.

**Figure 4.**
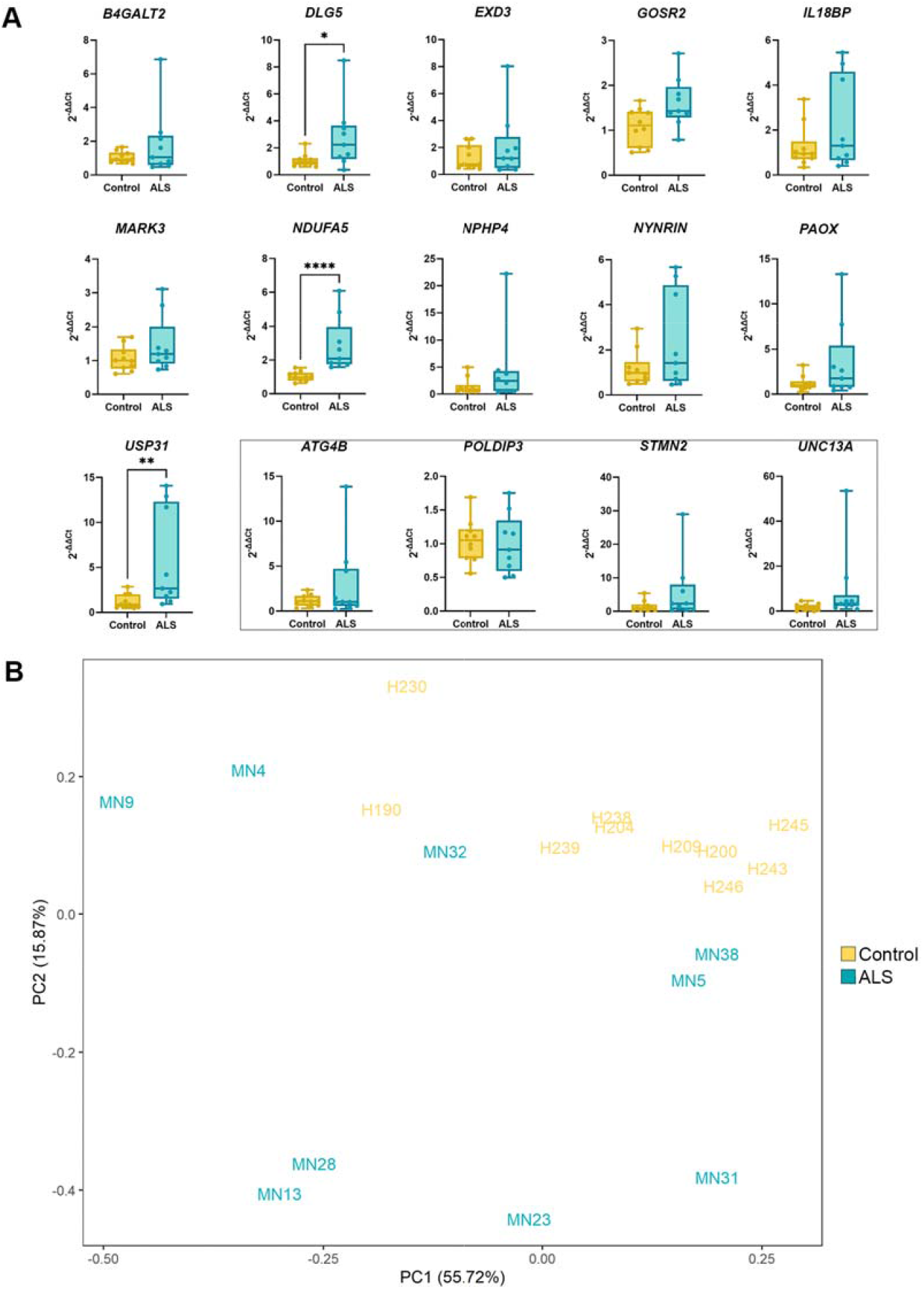
Differential exon usage in human ALS brain tissue. A) Fold changes in DEU events between ALS (n= 9, blue) and control (n= 10, yellow) motor cortex tissue by qPCR (Mann-Whitney U test: *, p = 0.035; **, p = 0.006; ****, p <0.0001). B) Principal component analysis of the ΔCts from all DEU events for each case.

When the ΔCts in each DEU event of the panel were considered together to represent a DEU/cryptic exon ‘profile’, principal component analysis (PCA) showed tight clustering of control cases, while ALS cases were more variable (**Fig. 4B**). Two control cases that clustered slightly apart from other controls (H230 and H190) had age-related changes indicated in their pathology reports (**Table S2**). Along the PC1 axis, MN4, MN9, MN13 and MN28 separated clearly from controls, while along the PC2 axis, MN5, MN13, MN23, MN28, MN31 and MN38 separated from controls, suggesting that TDP-43 LOF DEU/cryptic exon profiles differ between individual ALS cases but that TDP-43 LOF occurs in ALS brain tissue.

## Discussion

The heterogeneity in disease pathogenesis between individuals is one of the primary hurdles in ALS research. Determining common mechanisms of disease is challenging yet essential for developing therapeutics. Even with various genetic and environmental factors at play, the known common denominator in most ALS/FTD is TDP-43 proteinopathy (1, 2, 45). It is therefore critical to understand whether TDP-43 proteinopathy induces loss of its RNA processing function, and to identify specific gene changes caused by any such loss of function, to monitor and target the pathomechanisms that affect the largest patient population. Here we modelled TDP-43 LOF, which is caused by the interdependent processes of TDP-43 nuclear clearance (5, 46) and TDP-43 aggregation (2, 47). TDP-43 functions include splicing regulation and cryptic exon repression (9, 10) upon which the present study focuses. In testing the effects of TDP-43 LOF, we have identified key splicing events under regulatory control by TDP-43 and shown that changes in these splicing events are detectable in human brain tissue and may also be amenable to sampling in peripheral tissues.

Previous studies of TDP-43 splicing function indicated that a loss of TDP-43 function does not require its aggregation (8, 46), and that those splicing events are species and cell-type specific. For instance, cryptic exons resulting from TDP-43 depletion in a human-derived cell line did not concur with those in mouse embryonic stem cells, although mouse cells also underwent splicing changes (11, 41). We too observed species-specific splicing changes in our meta-analysis of transcriptomes following TDP-43 knockdown in various cell types (4). Further, within human models, there are cell-type-independent and cell-type-specific TDP-43 splicing targets (4), including in non-neuronal cell types such as Schwann cells and astrocytes that do not typically exhibit TDP-43 aggregation (48, 49). It is increasingly clear that identifying TDP-43 splicing targets in human disease requires human model systems, and that such targets may reveal TDP-43 dysfunction in neurons *and* other cell types even in the absence of TDP-43 aggregation.

In this study we employed non-neuronal cultures of human brain origin to interrogate TDP-43 splicing targets: primary human post-mortem brain pericytes with TDP-43 silenced by siRNA. In the present study, human brain pericytes were directly isolated from post-mortem brain tissue of multiple donors without reprogramming or transformation, thus incorporating donor-specific polymorphisms and retaining epigenetic signatures of ageing. Indeed, we found that human brain pericytes with TDP-43 depletion reliably recapitulated cryptic exon targets identified in previous studies in *POLDIP3, ATG4B*, and *UNC13A* (**Fig. 2**), even though the expression of *UNC13A* is mainly neuronal (50-52).

The top changed DEU/cryptic exons identified following TDP-43 knockdown in pericytes occurred in the genes *EXD3, DLG5, PAOX, NPHP4, B4GALT2* and *NYNRIN*, all of which were increased with TDP-43 knockdown. Events in *EXD3, DLG5, B4GALT2* and *NYNRIN* had also been detectable in RNA-Seq data from other human TDP-43 LOF models and/or TDP-negative neuronal nuclei from ALS/FTD tissue (4). However, only the cryptic exon in *DLG5* was significantly increased in ALS motor cortex tissue compared to controls in the present study. Brain tissue comprises various cell types, and both brain tissue sampling and disease can alter the relative proportions of cell types between samples. This variability will render DEU/cryptic exons difficult to detect if they only occur in a proportion of transcripts of that gene, a proportion of cells of a given cell type, or in rare cell types such as pericytes. Single-cell quantitative approaches such as single-cell RNA sequencing, RNAScope or high-throughput *in situ* hybridisation methods (53) may be used in future to discriminate between these possibilities for DEU events identified in cultured pericytes. However, these splicing changes are reproducible across many TDP-43 LOF models and will report upon TDP-43 splicing function in suitable assays if not in bulk tissue.

TDP-43-regulated DEU events that were significantly altered in ALS brain tissue included exonic elements in the genes *DLG5, NDUFA5* and *USP31* (**Fig. 4A**); evidence that TDP-43 LOF is at least partially detectable in ALS tissue. DLG5 is a scaffold protein involved in cell polarity, cytoskeleton maintenance (54), microtubule stability (55) and synaptogenesis (56); NDUFA5 is part of complex I of the mitochondrial respiratory chain (57); and USP31 is involved in deubiquitylation (58). The functions of these genes may be significant to pathomechanisms of disease. Tam and colleagues (59) analysed the transcriptomes of 148 ALS postmortem cortex samples and found that the majority (61%) displayed hallmarks of proteotoxic and oxidative stress, to which USP31 and NDUFA5 may contribute, given that deubiquitinating enzymes fine-tune protein degradation (USP31 function) (60) and oxidative phosphorylation produces reactive oxygen species (NDUFA5 function) (61). The function of DLG5 may have parallels with other ALS-related genes involved in cytoskeletal dynamics such as *DCNT1* (62), *PFN1* (63, 64), *TUBA4A* (65) and *STMN2* (17, 19).

Based on the genomic location of the DEU events, cryptic exon/retained intron events occur downstream of exon 31 for *DLG5* and exon 13 for *USP31*, while the exon usage change occurs at the 3’UTR for *NDUFA5*. The emergence of cryptic exons/retained introns may cause nonsense-mediated decay or altered translation, and 3’UTR alterations can result in a plethora of dysregulation possibilities, including altered mRNA stability, localization, translation or interaction with other RNA-binding proteins (66-70). The effects of these DEU events on the overall function of the genes in which they occur warrant further investigation. In addition to showing that TDP-43 LOF occurs in ALS tissue at the transcriptomic level, it will also be essential to determine cognate protein levels in ALS tissue to determine the translational and functional effect of TDP-43 LOF.

The variability in magnitude of individual DEU/cryptic exons and the spread of combined DEU/cryptic exon ‘profiles’ by PCA between brain tissues from different individuals suggests that heterogeneity in ALS may extend to TDP-43-regulated splicing changes. It is likely that the heterogeneity in DEU/cryptic exon expression in tissue is underpinned by the same factors, such as regional TDP-43 aggregate load (71) and genetic causes/ modifiers (72, 73), that drive clinical heterogeneity of ALS.

Some DEU events have been proposed to have biomarker potential, such as a cryptic exon within *ATG4B* that emerged with TDP-43 LOF and in ALS (11, 43) as well as in Alzheimer’s disease with TDP-43 pathology (46, 74). Diagnostic approaches include testing DEU changes or cryptic exon emergence with PCR or RNA-sequencing of biofluids (75, 76). Another emerging approach is designing aptamers or antibodies to cryptic peptides or neoepitopes that result from DEU events (77, 78). Indeed, the isoform switch of POLDIP3 from variant-1 to variant-2 due to exon skipping in *POLDIP3* has also been proposed to be a measurable protein biomarker due to variant-2 being rarely detected in normal tissues (44). Of our panel, *POLDIP3, ATG4B* and *MARK3* are the most ubiquitously expressed, with relatively strong expression in both brain and whole blood. We contend that DEU/cryptic exon events should occur in both brain and in tissue accessible during life, such as blood, cerebral spinal fluid (CSF) or fibroblasts, to act as useful biomarkers of the dysfunction caused by TDP-43 LOF. Further, specific profiles of DEU/cryptic exon events may contribute to, and/or report on, the heterogeneity in ALS. If individual patient profiles using DEU/cryptic exon panels such as the one presented here are shown to predict treatment effect, such panels may be beneficial for stratifying ALS/FTD cases for personalised treatment.

## Conclusion

A subset of DEU/cryptic exon events that occur in TDP-43 LOF models are detectable in ALS/FTD brain tissue, validating TDP-43 LOF as a disease mechanism. However, the magnitude of change of a given DEU/cryptic exon event varies between individuals. Further, because DEU/ cryptic exon events are enriched in specific cell types they are not always detectable in tissues of mixed cell type. TDP-43 LOF DEU/cryptic exon events will be most useful as biomarkers if they also occur in readily accessible tissues such as blood, CSF, or fibroblasts.

## Methods

### Donor pericyte line isolation, culture and transfection

All protocols involving human-derived samples were approved by the University of Auckland Human Participants Ethics Committee, and all methods were carried out in accordance with the approved guidelines. Post-mortem, non-neurodegenerative disease autopsy human brain tissue from 10 donors (7 male, 3 female; mean age 66.6 ± 13.5 years (SD)) was obtained from the Neurological Foundation Human Brain Bank and Hugh Green Biobank (University of Auckland, New Zealand) (**Table S1**). Tissue from the middle temporal gyrus (MTG) and motor cortex (MC) was processed for the isolation of mixed glial cultures, with early passages (up to passage 5) containing a mixture of pericytes, astrocytes and microglia, as previously described (79). Cells were maintained as described previously (42, 80). Due to the proliferation of pericytes, which overgrow the non-proliferative glial cells, later passage cultures contain almost exclusively pericytes (81-83). To examine TDP-43 loss of function (LOF) in pericytes, cells were transfected with scrambled (scr) small interfering RNA (siRNA) or siRNA against *TARDBP* (siTDP) as described previously (42).

### Donor human brain tissue

To validate whether splicing changes in pericytes with TDP-43 LOF were detectable in ALS, donor human brain tissue from control and ALS cases was examined. Cases were matched as closely as possible (**Table S2**). Control cases (6 male, 4 female; mean age 65.4 ± 11.4 years (SD)) were included in this study based on the absence of neurological conditions or significant neuropathologies. Age-related pathological changes were deemed acceptable. ALS cases (4 male, 5 female; mean age 62.2 ± 13.8 years (SD)) were included in this study based on a clinical diagnosis of ALS by neurologists at Auckland City or Middlemore Hospitals (New Zealand) during life. ALS pathology was confirmed by neuropathologists at Auckland City Hospital based on the detection of phosphorylated TDP-43 in the motor cortex and/or spinal cord post-mortem. The left hemispheres of donated brains obtained at autopsy were immediately dissected into functional blocks and frozen using CO_2_ powder and stored at -80 °C as described previously (84). Fresh-frozen motor cortex (MC) tissue (maximum input weight 20 mg) was used for RNA isolation.

### RNA isolation and RNA-sequencing

Following 96 h transfection with siRNA (scr or siTDP), cells plated at 15,000 cells/cm^2^ in 6-well plates were washed in ice-cold 1X phosphate-buffered saline (PBS) and RNA extraction and purification was performed using the RNAqueous®-Micro Total RNA isolation Kit (Ambion) as per manufacturer’s instructions. RNA isolation from fresh-frozen brain tissue was performed using the RNeasy® Mini kit as per manufacturer’s instructions. To homogenise the tissue, it was first disrupted in 600 μL lysis buffer (Buffer RLT) using pre-chilled pestles then passed through a 23-gauge needle 10 times. RNA integrity (RNA integrity number; RIN) was assessed using the Agilent RNA 6000 Pico Kit (Agilent Technologies) on an Agilent 2100 Bioanalyzer (Agilent Technologies). RNA from paired samples (scr and siTDP) from 9 donor cell lines (**Table S1**) were sent to AnnoRoad Gene Technology (Beijing, China) for quality check, library preparation and sequencing using the Illumina NovaSeq6000 platform and analysed as described previously (4, 42). Reads were aligned to the GRCh38 reference genome with HISAT2 (85) and quantified with StringTie (86). R package *DESeq2* (87) was used for differential expression analysis and *DEXSeq* (88) was used for exon usage analysis. RNA-seq coverage traces were viewed on Integrative Genome Viewer (IGV) (89).

### Quantitative reverse transcription PCR (qPCR) and primer design

To validate DEGs and DEU events detected by RNA-sequencing, RNA from three pairs of pericyte cases, considered a ‘replication cohort’ (**Table S1**), was tested by qPCR. Either 0.5 or 1 μg of RNA was used for cDNA synthesis. RNA was treated with DNase I (Promega) (1 μg DNase I/ 1 μg RNA) and cDNA was synthesised using a Superscript® First-Strand Synthesis IV kit (ThermoFisher Scientific). qPCR was performed with Platinum® SYBR® Green qPCR SuperMix-UDG with Rox (ThermoFisher Scientific) with a QuantStudio™ 12K Flex Real-Time PCR system (Applied Biosystems). The qPCR program was set to 50 °C for 2 min, 95 °C for 10 min, and 40 cycles of 95 °C for 15 s and 60 °C for 1 min. Relative quantification was determined using the delta-delta Ct method with normalization to the housekeeping gene *GAPDH*. The fold changes are reported using 2^-ΔΔCt^ values, with the ΔCt for each case calculated relative to the average ΔCt of all control cases.

Primers were designed using PrimerBlast and Geneious 10.2.6 software (90). PrimerBlast parameters were set to 80-120 bp amplicon length, maximum primer T_m_ difference 2 °C, and GC content 50%-60%. Maximum self-complementarity was set to 4 and self-3’ complementarity was set to 2. Primer sequences can be found in **Table S3**.

### Immunocytochemistry and image acquisition

To confirm the extent of TDP-43 silencing in human brain pericytes, following 96 h transfection with siRNA (scr or siTDP), cells plated at 15,000 cells/cm^2^ in 96-well plates were fixed in 4% PFA for 10 minutes and washed in 1X PBS containing 0.2% (v/v) Triton X-100 (PBS-T). Cells were incubated with rabbit polyclonal anti-TDP-43 (1078-2-2-AP, Proteintech) diluted 1:2000 in PBS-T with 1% goat serum overnight at 4 °C. Cells were then washed twice with 1X PBS and incubated with goat anti-rabbit AlexaFluor™ Plus 488 (A32731, ThermoFisher Scientific) diluted 1:500 in PBS-T with 1% goat serum overnight at 4 °C. Cells were then washed twice with 1X PBS and nuclei were counterstained with 20 nM Hoechst 33258 (Sigma-Aldrich) diluted in PBS-T for 10 minutes. Images of immuno-stained cells were acquired using an Image Xpress Micro XLS High Content Screening system (Molecular Devices) at 10X/0.3 NA Plan Fluor magnification using filters DAPI (excitation range 381-399 nm) and FITC (excitation range 474-496 nm).

### Publicly available data

Gene expression heatmap displaying genes with DEU events from TDP-43 LOF was obtained from the Genotype-Tissue Expression (GTEx) Portal V8 (www.gtexportal.org) on 09 January 2023 (dbGaP accession number phs000424.v9.p2) using the Multi-Gene Query.

### Statistical analysis

Statistical analysis for differential exon usage was performed by the R package *DEXSeq*, which used *DESeq2* Wald test and false discovery correction with Benjamini-Hochberg method (87, 88). Significant padj value was set to <0.05. Paired t-test was performed for scr versus siTDP pericyte comparisons, whilst Mann-Whitney U test was performed for control versus ALS brain tissue comparisons, using GraphPad Prism 9.0 software (La Jolla, CA, USA). Significant p value was set to <0.05. Principal component analysis (PCA) was performed for each control and ALS case based on the ΔCT values for each qPCR primer pair using the prcomp function within R software.

## List of abbreviations

ALS: Amyotrophic lateral sclerosis
BBB: Blood-brain barrier
BSCB: Blood-spinal cord barrier
cDNA: Complementary DNA
Ct: Cycle number threshold
DEU: Differential exon usage
FTD: Frontotemporal dementia
LOF: Loss of function
log2FC: Log2 fold change
MC: Motor cortex
mRNA: Messenger RNA
MTG: Middle temporal gyrus
PBS: Phosphate-buffered saline
PCA: Principal component analysis
qPCR: Quantitative polymerase chain reaction
RIN: RNA integrity number
RRM: RNA recognition motif
Scr: Scrambled siRNA
SD: Standard deviation
siRNA: Small interfering RNA
siTDP: siRNA against *TARDBP*
SNP: Single nucleotide polymorphism
TDP-43: TAR DNA binding protein of 43 kDa
UTR: Untranslated region

## Declarations

### Ethics approval and consent to participate

All protocols were approved by the University of Auckland Human Participants Ethics Committee (New Zealand) and carried out as per approved guidelines.

### Consent for publication

Informed consent was obtained from all tissue donors.

### Availability of data and materials

Raw and processed RNA-sequencing data is available on NCBI GEO datasets, series accession number GSE223747.

Interactive database is available online at https://www.scotterlab.auckland.ac.nz/research-themes/tdp43-lof-db-v2/.

## Competing interests

The authors declare that they have no competing interests

## Funding

This work was supported by grants from the Royal Society of NZ [RDF-15-UOA-003], Marcus Gerbich and Amelia Pais-Rodriguez, Motor Neurone Disease NZ, Freemasons New Zealand, Matteo de Nora, Coker Family Trust, Sir Thomas and Lady Duncan Trust, Health Research Council of New Zealand, Hugh Green Foundation and PaR NZ Golfing. No funding body played any role in the design of the study, nor in collection, analysis, or interpretation of data nor in writing the manuscript.

## Authors’ contributions

Experiments and analysis by MCC. Study design and manuscript draft by MCC and ELS. RNA extraction from human brain tissue by BR. RNA-seq trimming and alignment by JW. Coordination of banking, use of, and methods for culture of human pericytes via The Hugh Green Biobank by MD. Coordination of banking and use of human tissue via the Neurological Foundation Human Brain Bank by RLMF and MAC. All authors have read and approved the final manuscript.

## Acknowledgements

This publication is dedicated to the patients and families who contribute to our research. We thank Marika Eszes at the Centre for Brain Research, University of Auckland, New Zealand, and the Neurological Foundation of New Zealand for their ongoing financial support of the Human Brain Bank. The imaging data reported in this paper were obtained at the Biomedical Imaging Research Unit, operated by the Faculty of Medical and Health Sciences’ Technical Services at the University of Auckland.

## Notes

https://www.ncbi.nlm.nih.gov/geo/query/acc.cgi?acc=GSE223747

### Competing Interest Statement

The authors have declared no competing interest.

### Summary of Updates

Revisions include a link to a database that allows comparison of differentially expressed genes and exon usage changes between RNA-sequencing studies of TDP-43 loss of function (https://www.scotterlab.auckland.ac.nz/research-themes/tdp43-lof-db-v2/). Version 1 of this database (as published in Transcriptional targets of amyotrophic lateral sclerosis/frontotemporal dementia protein TDP-43 - meta-analysis and interactive graphical database. Disease Models & Mechanisms. 2022 Sep 1;15(9):dmm049418) is still available at https://www.scotterlab.auckland.ac.nz/research-themes/tdp43-lof-db/ The v2 database been updated to include the human brain-derived pericyte data of this study.

