## supplementary tables for "A panel of TDP-43-regulated splicing events verify loss of TDP-43 function in amyotrophic lateral sclerosis brain tissue"

**Supplementary data**

***Table S1: Pericyte cell line donor information***

| Case ID | Brain region | Age at death (y) | COD | Sex | PMD (h) | RIN  (scr/siTDP) |
| --- | --- | --- | --- | --- | --- | --- |
| H189 | MC | 41 | Asphyxia | M | 16 | 10.0/10.0 |
| H189 | MTG | 41 | Asphyxia | M | 16 | 10.0/9.8 |
| H194 | MC | 68 | Coronary atherosclerosis | M | 22.5 | 9.0/8.9 |
| H209 | MTG | 48 | Ischaemic heart disease | M | 23 | 9.8/10.0 |
| H227 | MC | 78 | Cerebrovascular accident | F | 4 | 9.6/8.7 |
| H238 | MTG | 63 | Dissecting aortic aneurysm | F | 16 | * |
| H239 | MTG | 64 | Ischaemic heart disease | M | 15.5 | * |
| H243 | MTG | 77 | Ischaemic heart disease/ coronary atherosclerosis | F | 13 | * |
| H244 | MC | 76 | Ischaemic heart disease/ coronary atherosclerosis | M | 16 | 9.8/9.5 |
| H245 | MC | 63 | Asphyxia | M | 20 | 9.9/7.2 |
| H246 | MC | 88 | Type II myocardial infarction | M | 17 | 10.0/10.0 |
| H246 | MTG | 88 | Type II myocardial infarction | M | 17 | 9.9/5.4 |

Abbreviations: MC, motor cortex; MTG, middle temporal gyrus; COD, cause of death; PMD, post-mortem delay; RIN: RNA integrity score**.**

* Replication cohort case for qPCR (not RNA-sequenced)

***Table S2: Brain tissue donor information***

| Case ID | Brain region | Age at death | Sex | PMD (h) | Pathology | RIN |
| --- | --- | --- | --- | --- | --- | --- |
| H190 | MC | 72 | F | 19 | Age-related microscopic changes | 5.7 |
| H200 | MC | 56 | M | 23 | No significant pathologies | 7.8 |
| H204 | MC | 66 | M | 9 | No significant pathologies | 6.5 |
| H209 | MC | 48 | M | 23 | No significant pathologies | 7.8 |
| H230 | MC | 57 | F | 32 | Age-related cerebral changes | 5.3 |
| H238 | MC | 63 | F | 16 | No significant pathologies | 5.2 |
| H239 | MC | 64 | M | 15.5 | No significant pathologies | 7.8 |
| H243 | MC | 77 | F | 13 | No significant pathologies | 6.3 |
| H245 | MC | 63 | M | 20 | No significant pathologies | 6.5 |
| H246 | MC | 88 | M | 17 | Atherosclerosis | 5.6 |
| MN4 | MC | 41 | M | 7 | ALS | 2.2 |
| MN5 | MC | 55 | F | 5 | ALS | 7.8 |
| MN9 | MC | 88 | M | 36 | ALS | 7.4 |
| MN13 | MC | 55 | M | 10 | ALS | 8 |
| MN23 | MC | 79 | F | 27 | ALS (*C9orf72*) | 5.8 |
| MN28 | MC | 62 | F | 14 | ALS (*C9orf72*), mild AD | 6.2 |
| MN31 | MC | 58 | F | 19 | ALS | 7.8 |
| MN32 | MC | 59 | M | 6 | ALS | 8.1 |
| MN38 | MC | 63 | F | Not recorded | ALS | 5.9 |

Abbreviations: MC, motor cortex; AD, Alzheimer’s disease; PMD, post-mortem delay; RIN: RNA integrity score**.**

***Table S3: Primers for qPCR***

| Gene name | Forward primer  (5’ to 3’) | Reverse primer  (5’ to 3’) | Genomic location of amplicon (GRCh38) |
| --- | --- | --- | --- |
| *ATG4B* | TCATAACCTGCCCCAATCCC | CCTTGATTCGCCCACGAGAT | Chr2: 241,670,415 – 214,670,495 |
| *B4GALT2* | GGGTGTCCCTGTGATTCTGT | GCTGTGGTTCCAAGAGAGCG | Chr1: 43,980,138 – 43,980,228 |
| *DLG5* | ACTTGTCAGGAACGTGCTCA | AAGGCTGTTTTCTTAGCCGTGG | Chr10: 77,793,847 – 77,793,967 |
| *EXD3* | CTCCAGTGAGACAACCCTG | GCAAGCTGAAAGGTCTGTGA | Chr9: 137,365,318 – 137,365,415 |
| *GAPDH* | CATGAGAAGTATGACAACAGCCT | AGTCCTTCCACGATACCAAAGT | Chr12: 6,537,181 – 6,537,386 |
| *GOSR2* | GCCCTGATGGGAAGGTCTGT | GAAAGGCCACATCACCTTGGA | Chr17: 46,964,876 – 46,964,961 |
| *IL18BP* | TTGCGATACTAGCCTGCACC | GGCACAGCCAGCAACTGATA | Chr11: 72,003,573 – 72,003,683 |
| *MARK3* | GATGTGCAACCTATGCGCC | TCACACACATACCGTCCACTC | Chr14: 103,503,407 – 103,503,526 |
| *MMAB* | TCAGTCCAACCAGACAGGGA | GATGGGGGTGCAGCTTAGAG | Chr12: 109,556,293 – 109,556,397 |
| *NDUFA5* | CAAACACCAGTAAGGTCCCC | TTTTCAGTCCCGTATCTGCC | Chr7: 123,540,313 – 123,540,412 |
| *NPHP4* | ACCCTTCCCTTGTGGGTCT | GCAGTGATCGTATGCAGTGT | Chr1: 5,888,462 – 5,888,549 |
| *NYNRIN* | CTGGTCCTGGGTTGGAGTTG | CCTGGCATGAGAGTTGCCTA | Chr14: 24,412,035 – 24,412,106 |
| *PAOX* | CGTGAGCCCTTTTCTCGTCC | GTCTGGGCAAGGAAACACAGA | Chr10: 133,391,160 – 133,391,262 |
| *POLDIP3* | GTTCCACAGCAGAAAGCCAT | CTGTTTGGCCTGGTGGTTAT | Chr22: 42,601,969 – 42,602,056 |
| *STMN2* | TGTGTGAGCATGTGTGCGT | CCACAAGCCGCATTCACAT | Chr8: 79,616,873 – 79,616,955 |
| *TARDBP* | GGGGAAATCTGGTGTATGTTG | CGGATGTTTTCTGGACTGCT | Chr1: 11,013,929 – 11,016,919 |
| *UNC13A* | GGGACCTTGGGATGGTTTCA | AGCACAAAAGCTCCCTTCCA | Chr19: 17,628,010 – 17,628,097 |
| *USP31* | CAAAACCGTAAATGGGGAGCG | CTGACTGATTGGGGAGTGGT | Chr16: 23,079,637 – 23,079,735 |
